# Cryptic Insect Microbiome Compositions Unveiled with Full-Length 16S Sequencing

**DOI:** 10.64898/2026.01.22.701149

**Authors:** Charles J. Mason, Thorsten E. Hansen, Nicholas Newell, Scott M. Geib

## Abstract

Insects have interconnected relationships with gut microorganisms which can have impacts on behavior and survival. Understanding foundational ecological principles of the insect-associated microorganisms is important to understand how insects utilize microbes to cope with stress and different environments. Common microbiome surveys of insect microbiomes utilize 16S rRNA sequencing of metagenomic DNA. For amplicon-based surveys of insect microbiome, fragment length and 16S rRNA subunit choice may have unintended biases, and some primer combinations may under-represent genus or species richness. Contemporary solutions target sequencing of the entire 16S rRNA fragment. Here, we illustrate the benefit of full-length 16S rRNA sequencing in improving sample resolution compared to V4 which provides new insight into the gut microbiome community composition of an invasive insect. We evaluated the gut microbiome of mass-reared medfly males (Mediterranean fruit fly, *Ceratitis capitata*) that were collected across a nine-month sampling period. Full-length 16S rRNA PCR products of samples were prepared into a Kinnex 16S rRNA library, sequenced on a PacBio Revio system, and the resulting HiFi amplicon data was processed into amplicon sequence variants (ASVs). Our findings reveal substantial differences in bacterial compositions across different medfly cohorts when sampling the full-length amplicons which were not detectible when only considering the V4 regions. Strain-level gut microbiome variation were supported with genomes assembled from shotgun metagenomic sequencing. Our findings support that long-read sequencing of full-length 16S rRNA amplicon uncover ecologically important interactions between host and gut microbiomes and serve as a bridge between short-fragment sequencing and shotgun metagenomics.

## Introduction

Plant-feeding insects harbor gut microbial assemblages that mediate important ecological interactions [1, 2]. The contributions that gut bacteria have on digestion, detoxification, and immunity can be leveraged by herbivore hosts to have broader impacts on lifespan and reproduction. Knowledge of the ecology and evolution of insect microbiomes can provide valuable insight into species distribution, host plant usage, and pest management [3-5].

Foundational exploration of insect gut microbial ecology has most often relied on short fragment 16S ribosomal RNA gene (16S) amplicon sequencing using high-throughput paired-end short read sequencing. While these approaches are excellent tools and provide valuable insight into the microbial ecology of insects, for certain taxonomic groups, the region of the 16S may inadvertently bin distinct species or strains into the same variant or taxonomic unit [6-8]. This can be problematic for plant-feeding insects who are often associated with Enterobacteriaceae [4, 9], where even at the genus level isolates can be difficult to delineate with 16S [10-13] and short fragments may exacerbate these issues [14]. 16S amplicon sequence lengths are limited in part by the sequencing technology, with sequencing by synthesis chemistries not able to effectively sequence larger amplicons [8]. Shotgun metagenomics and metagenomic-assisted assemblies provide obvious advantages over 16S amplicons for delineating strains and species [15, 16], but cost, analytical complexity, and host DNA interference my limit the ability to routinely screen larger sample sizes efficiently. Long-read approaches encompassing the all 16S hypervariable regions, especially with high-quality reads, aim to overcome these limitations to facilitate more detailed insights into ecological underpinnings of gut microbiomes of closely related microbial taxa with an advantage of allowing for greater sample sizes and improved statistical inferences [17-19]. Better differentiation of bacterial species or strains within an insect’s gut microbial community may provide improved insight into dynamics of microbiota in the face of perturbations and their impacts on the host.

In this study, we took advantage of samples from a study exploring cohort variation in bacterial communities of the invasive tephritid fruit fly, the Mediterranean fruit fly (medfly; *Ceratitis capitata*). In our prior study, samples were sequenced targeting the 16S V4 hypervariable region using 300 base-pair paired-end techniques [20]. A pertinent observation from those experiments were that there were few detectable differences in the gut microbiota across time and treatments and that the systems were largely comprised of ASVs classified as Enterobacteriaceae (genus *Klebsiella*). Based on separate experiments we conducted in medfly tracking a specific *Enterobacter* isolate and limitations observed with the V4 hypervariable region [21], we hypothesized that increased sequence lengths may reveal greater bacterial biodiversity than what was initially observed with partial 16S reads. To test this, here, we performed near full-length 16S sequencing using PacBio Kinnex, a MAS-seq method involving read concatenation [22], to revisit these specimens to determine if complete surveying of the 16S rRNA amplicons illuminates ecologically relevant aspects of the medfly gut microbiome. We then further interrogated and supported these findings by conducting shotgun metagenomic sequencing and co-assembled genomes with a subset of these samples.

## Methods

### Sample Origin

Sample collection occurred over of a nine-month period in 2023 [20]. Medfly males were mass-reared by the California Department of Agriculture (CDFA) in Waimanalo, HI, USA and shipped to the USDA ARS Daniel K. Inouye Pacific Basin Agricultural Research Center in Hilo, HI, USA. A treatment involving sterilization of males with gamma irradiation was a component of the original study design. Samples sizes were initially 10-11 individuals for each cohort and treatment upon initial processing. Extraction, pooling, and sequencing yielded at least eight samples for each combination. Negative control wells and positive controls (Zymo Research, Irvine, CA, USA) were sequenced along with samples.

### 16S Kinnex Amplification, Sequencing, and Processing

Samples were amplified using primers designed for PacBio Kinnex PCR (Pacific Biosciences, Menlo Park, CA, USA). Reactions were performed in 15 µL volumes with 0.25 µM of 27F and 1492R primers containing barcodes and nucleotides to enable Kinnex PCR. Reactions were performed using intercalating dyes on the iconPCR instrument (n6 Tec, Pleasanton, CA, USA), which allows for individual sample control such that amplification cycles may be ceased following a change in fluorescence so samples are not overamplified and are normalized [23, 24]. The iconPCR instrument was operated under ‘Slope mode’ to generate amplicons where PCR cycles were terminated in the exponential phase. Amplification was performed using Q5 Hi-Fidelity Hot Start Polymerase (New England Biolabs, Ipswich, MA, USA) supplemented with 1× EvaGreen (Biotium, Fremont, CA, USA). Thermocycler conditions were: 98°Cfor 30s, cycles of 98°Cfor 10s, 55°Cfor 15s, 72°Cfor 2 min, and a final extension of 72°Cfor 10 min. Number of actual cycles varied between samples with 30 cycles being the prescribed maximum (Supplemental Table 1).

Autonormalized PCR products were pooled in equal volumes (5 µL) following amplification and concatenated amplicons were produced according to the Kinnex 16S rRNA kit (Pacific Biosciences, Menlo Park). A single Kinnex library containing all individually barcoded samples was sequenced on a Revio 24M SMRT cell on a Pacific Biosciences Revio sequencer. Desegmentation of the Kinnex array into subcomponent amplicons and demultiplexing of amplicons to the sample level were conducted using SMRTLink v.13 software resulting in automated generation of per sample amplicon data files. For each sample, HiFi reads were pre-processed to retain reads with quality scores >Q30, trim primer sequences, and filter for final amplicon length of 1200-1500 nt reoriented on the positive strand [25, 26] prior to downstream analysis. Pre-processed data were then used to generate amplicon sequence variants (ASVs) using the DADA2 PacBio denoising algorithm using binned Revio quality scores [27, 28] in R v.4.4 [29]. Taxonomic designations of denoised ASVs were classified using the Naïve Bayesian Classifier with using Ribosomal Database Project Database v19 [30], to match our prior publication [21]. QA/QC metrics for each sample are provided in Supplemental Table 1.

### 16S Statistical Analysis

Statistical analyses were performed using R v.4.4.1 [29]. Amplicon samples had relatively even sampling depths and were rarefied to 3000 reads, which was selected to include >90% of test samples, and were greater than the number of reads in negative control wells and allowed for sample accumulation curves to saturate (Supplemental Figure 1). DNA controls produced sequences in proportions as anticipated (Supplemental Table 2). Rarefaction was performed with the package vegan v.2.6-8 [31] and the resulting data table was used for subsequent analyses. We computed Bray-Curtis dissimilarities between samples with vegan and performed ordination with principal coordinates analyses (PCoA). Permutational multivariate analysis of variance (PERMANOVA) and pairwise PERMANOVA [32] were conducted with cohort and treatment as explanatory variables. Alpha-diversity metrics were computed with vegan. Taxonomy bar plots were constructed at the genus level to illustrate taxonomic variability between specimens. A heatmap of the most prevalent ASVs (top 50) was constructed to evaluate patterns between cohorts with the package pheatmap v.1.0.12 [33], where ASVs were clustered with Manhattan distances using UPGMA.

We further evaluated 16S ASVs by constructing co-occurrence networks and delineating community modules across all cohorts and treatments. ASV-level associations were inferred using SPIEC-EASI v.1.1.1 [34], which accounts for compositionality and sparsity of microbiome datasets. SPIEC-EASI was performed using a neighborhood selection method [35] to infer covariance matrices using 99 permutations. ASV co-occurrence networks were constructed from association values computed by SPIEC-EASI in igraph v.2.1.4 [36]. Network modules were selected by the Leiden algorithm [37], a community detection algorithm that extracts communities from networks. The analysis was performed using ‘cluster_leiden’ with n_iterations = 50 with an initial_membership = 1 implemented in the igraph package. Module selection was performed with no assumptions of the number of community members. Networks were visualized using ggnet v.0.1.0 [38]. After selecting community modules, we evaluated if the relative abundances of ASVs assigned to those modules might help explain the survival of medfly from those same cohorts [20]. Relative abundances of each community module were summed for each cohort. We then performed correlations using Spearman coefficients between module mean relative abundance and cohort mortality metrics using rstatix v.0.7.2 [39]. Three metrics from the initial study were compared: percent mortality of flies 48h following dietary and water deprivation, percent mortality at fifteen days following eclosion, and median mortality (in days) of flies fed sucrose.

### Shotgun Metagenomic Sequencing and Processing

Following analysis of 16S data, five non-irradiated medfly samples from each cohort were sequenced for metagenomic assemblies. Samples (20 µL) were sheared (30 cycles; 30s on – 30s off) using a Biorupter Pico (Hologic Diagenode, Denvile, NJ, USA) before preparing libraries using a NEBNext Ultra II DNA Library Prep Kit. Libraries were sequenced on two cells on an Element AVITI sequencer using an AVITI 2×150 Sequencing Kit Cloudbreak FS High Output Kit (Element Biosciences, San Diego, CA, USA).

Read quality was assessed with FastQC v.0.11.9 [40] and Trimmomatic v.0.39 [41] was used to remove low quality reads and adapters. Host read removal, taxonomic profiling, contig assembly, assembled genome binning and bin refinement were performed using nfcore/mag v.4.0.0 [42] implemented in Nextflow v.25.04.2 [43] and singularity CE v.1.3.1 [44]. In the pipeline, fastp v.0.23.4 [45] was used for adapter removal and FastQC for further quality control. Reads mapped to PhiX (GCA_002596845.1, ASM259684v1) or the host medfly genome assembly (GCA_000347755.4) with Bowtie2 v2.4.2 were identified and removed [46].

High-quality paired end reads were then assembled into eight separate co-assemblies, based on their respective cohorts, using MEGAHIT v.1.2.9 [47] with parameters: --k-min 21, --k-max 141, --k-step 10, --min-count 3, --min-contig-len 500, and --no-mercy. The quality of these assemblies was assessed using QUAST v.5.0.2 [48]. The eight metagenome assemblies were then separately binned using MaxBin2 v.2.2.7 [49], MetaBAT2 v.2.15 [50], and CONCOCT v1.1.0 [51] to recover metagenome-assembled genomes (MAGs). The predicted bins across each cohort were combined using the DAS Tool v.1.1.6 [52]. MAG quality and completeness was determined with BUSCO v.5.7.1 [53, 54] using 124 universal single-copy orthologs from OrthoDB v.10 [55]. CheckM2 v.1.0.1 [56] was used to assess overall MAG completeness and contamination using universally trained machine learning modules. MAG taxonomy were assigned using GTDB-Tk v.2.4.0 [57]. FastANI v.1.34 [58] was used to compare the whole-genome average nucleotide identity (ANI) between the MAGs. Finally, SynTracker v.1.4.0 [59] was used to determine the biological relatedness of conspecific strains using genome synteny.

Taxonomic profiling based on 16S was performed on the metagenomic samples. Kraken2 v.2.14 [60] was used for taxonomic identification, Bracken v.3.1 [61] for abundance estimation, and KrakenTools v.1.2.1 [60] for analysis and data manipulation of output files. Two pre-built databases were used for these analyses. First, the BLAST Core Nucleotide Database (core_nt) was used to search for all possible k-mer gene matches to assign taxonomy in the metagenomic samples. Then the Silva v.138 16S database was used to identify taxonomy of bacteria in our metagenomic samples based off the traditional marker gene 16S.

### Shotgun Metagenomic Statistical Analysis

Metagenomic taxonomic data at species-level were transformed using cumulative sum scaling [62] before computing Bray-Curtis dissimilarities in phyloseq v.1.52.0 [63]. Comparisons between cohorts with these dissimilarities were conducted using PERMANOVA and pairwise PERMANOVA. Two metagenomic samples were culled due to low sequence representation (Cohort 071223) and an erroneously sequenced sample (Cohort 080923).

## Results

Near full-length 16S sequences revealed community-level differences associated with the microbiome of mass reared medfly (Figure 1). Ordination of ASVs by Bray-Curtis dissimilarities revealed a partitioning of samples in two-dimensional space with samples clustering by cohort (Fig 1A). PERMANOVA analysis identified statistical effects of cohort on ASV composition (R^2^ = 0.43; F_7,164_=17.5; p<0.001; Supplemental Table 3). Within this study, there was also a comparison between medfly sterilized with gamma radiation compared to those which were not. Gamma irradiation had marginally statistically significant effects on gut microbiota (R^2^ = 0.01; F_1,164_=1.96; p=0.069), but there was an interaction with cohort (R^2^ = 0.044; F_7,164_=1.80; p=0.003). Pairwise comparisons indicated that only one cohort (C062923; Supplemental Table 4) had statistically significant differences between irradiated and control flies. ASV richness and Shannon metrics were significantly different between cohorts (both p<0.001), but not with irradiation treatment or its interactions with cohort (Supplemental Figure 2). 16S sequences were primarily classified within the Enterobacteriaceae (Fig 1B), as *Enterobacter* as the most prevalent in the dataset. Heatmaps of individual ASVs illustrated multi-copy variation of the 16S in Enterobacteriaceae genomes (Fig 1C). Other bacterial taxa were represented, but at lower relative abundances (<2% across the dataset). Some ASVs were present across samples through time in the distinct cohorts, suggesting there are potentially common sources of the bacterial members.

**Figure 1:**
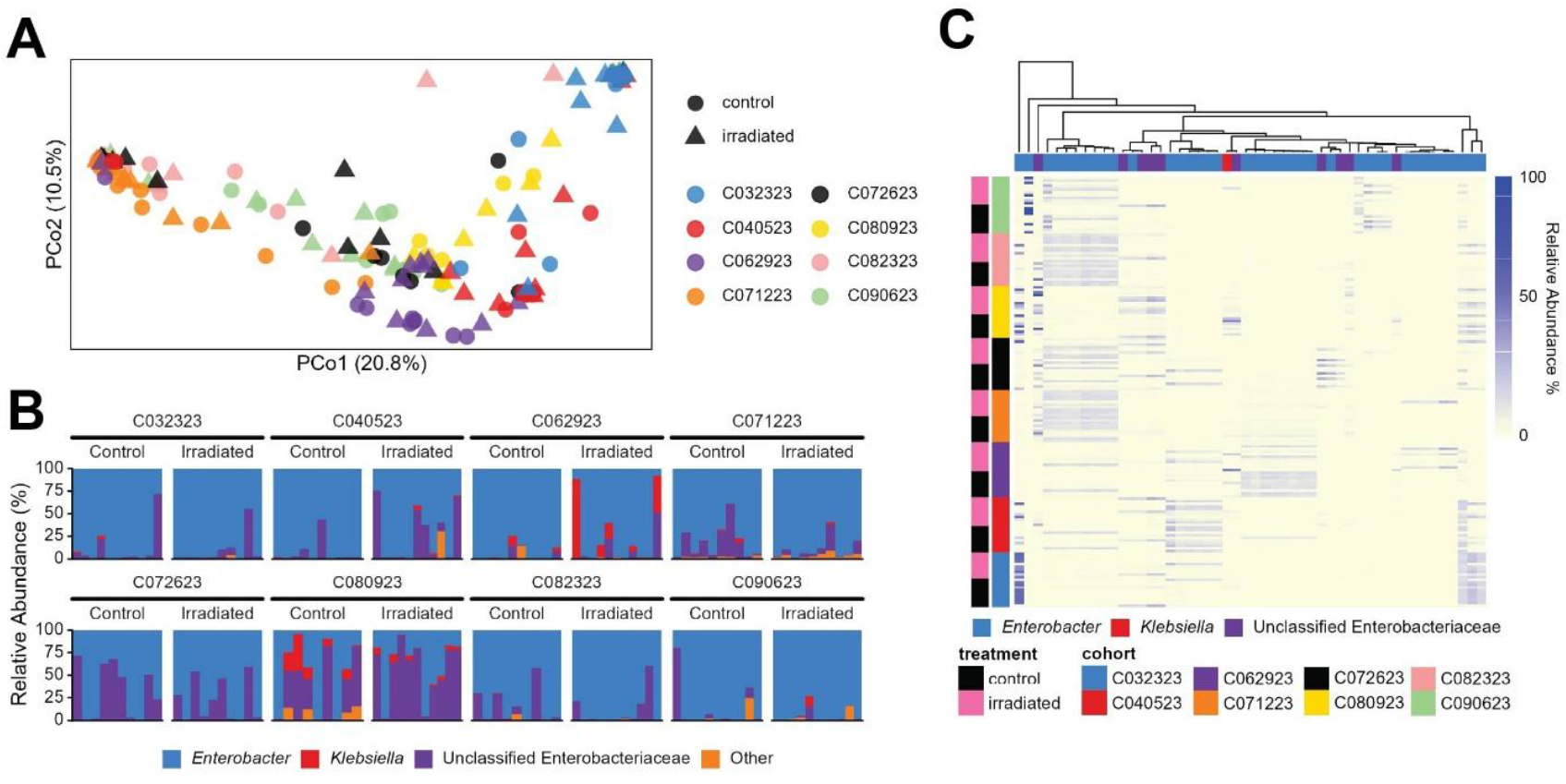
Impact of cohort and irradiation treatments on mass-reared medfly microbiome. (A) Principal coordinate analysis (PCo) of Bray-Curtis distances with full-length 16S rRNA ASVs. (B) Genus-level relative abundances (%) of ASVs in irradiated and non-irradiated medfly across eight timepoints. (B) Heatmap illustrating individual ASVs and taxonomic representation across samples. Clustered heatmaps by samples are illustrated in Supplemental Figure 3. ‘Other’ are sequences comprising <2% of the relative abundance across the total dataset.

Network analysis and module selection placed 261 16S ASVs into 18 community modules (Fig 2A; Supplemental Table 5). Module 1 had the highest representation, with the most ASVs belonging to this community type and highest relative abundance across different cohorts. (Fig 2B). Other modules exhibited more variable incidences across the eight cohorts. Mean relative abundances of modules for each cohort had relationships with fly survival across cohorts (Table 1). However, for time to 50% mortality of the cohort, relative abundances of Module 3 (ρ=0.54, p=0.029) had positive correlations with fly survival while Module 6 were negative (ρ= -0.59, p=0.016).

**Table 1:**
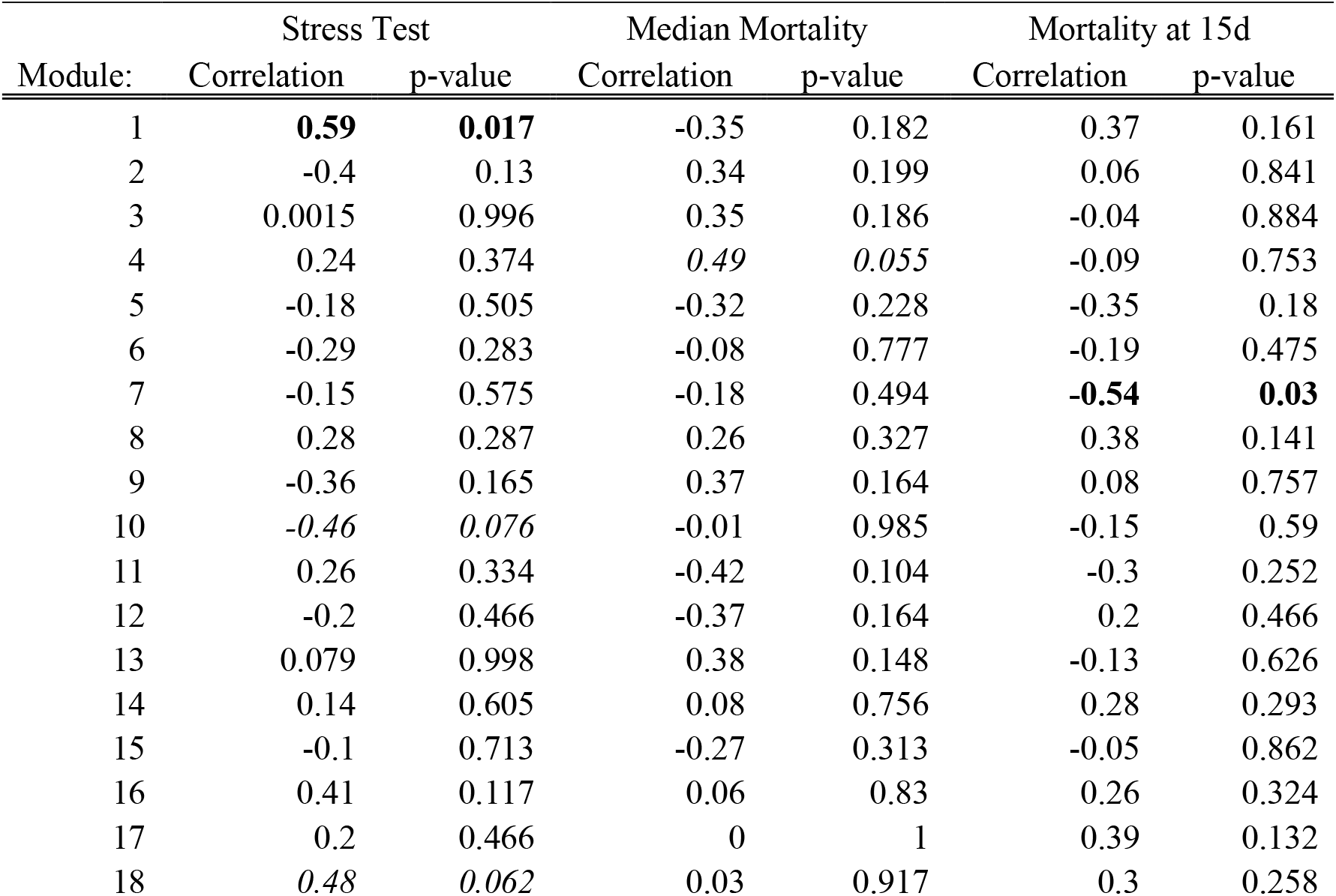
Spearman rank correlation coefficients between the mean relative abundance of ASVs in modules for each cohort with mortality measures of separate medflies from those cohorts. Stress Test - living flies following deprivation of food and water at 48h; Median mortality - median mortality in days for flies reared until death; Mortality at 15 and 30d - percent of flies alive at 15 and 30d after emergence. Bold values indicate statistical significance while italics represent marginally significant values (p < 0.1).

**Figure 2:**
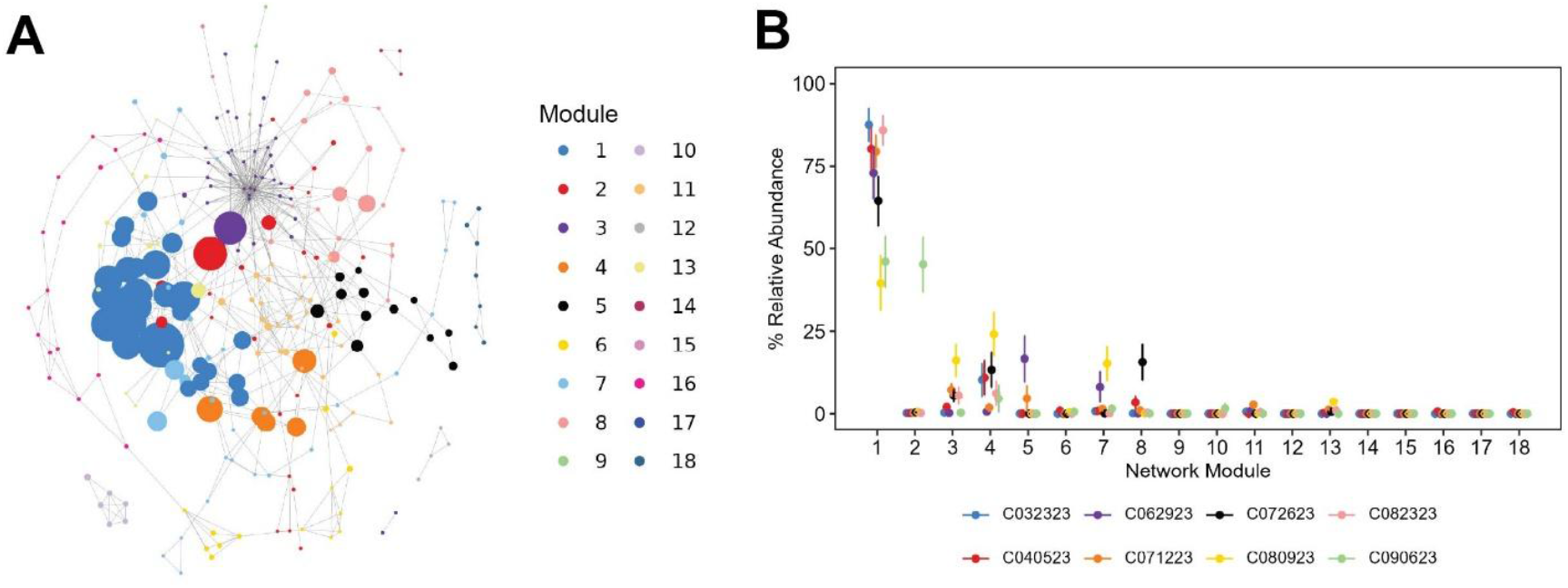
ASV co-occurrence network and community modules across all samples in dataset. (A) Network analysis where each point represents a distinct ASV and the size represents the relative abundance of that ASV of all samples. Community modules were detected using the Leiden algorithm. (B) Relative abundances (%) of ASVs within each community module and how they are distributed across different medfly cohorts (mean ± std error). ASVs within each module are provided in Supplemental Table 5.

Shotgun sequencing of a subset of samples revealed statistically significant differences in taxonomic makeup using Kraken2 inferred taxonomy (R^2^ = 0.35; F_7,37_=2.26; p=0.005; Figure 3A). While there was not a significant difference between all cohorts, there was divergence between a subset (Supplemental Table 6). Like full-length 16S data, the *Enterobacter* dominated at the genus level (Figure 3B). The assembled MAGs from the different cohorts generated complete *Enterobacter* genomes from seven of the eight cohorts (Figure 3C, Supplemental Tables 6-8). Assembled genomes included three *E. roggenkampii*, two *E. dykesii*, one *E. mori*, and one *E. soli*. A synteny-based analysis within species indicated that there was a mismatch between assemblies, which suggests different strains within each cohort (Supplemental Table 9 &10).

**Figure 3:**
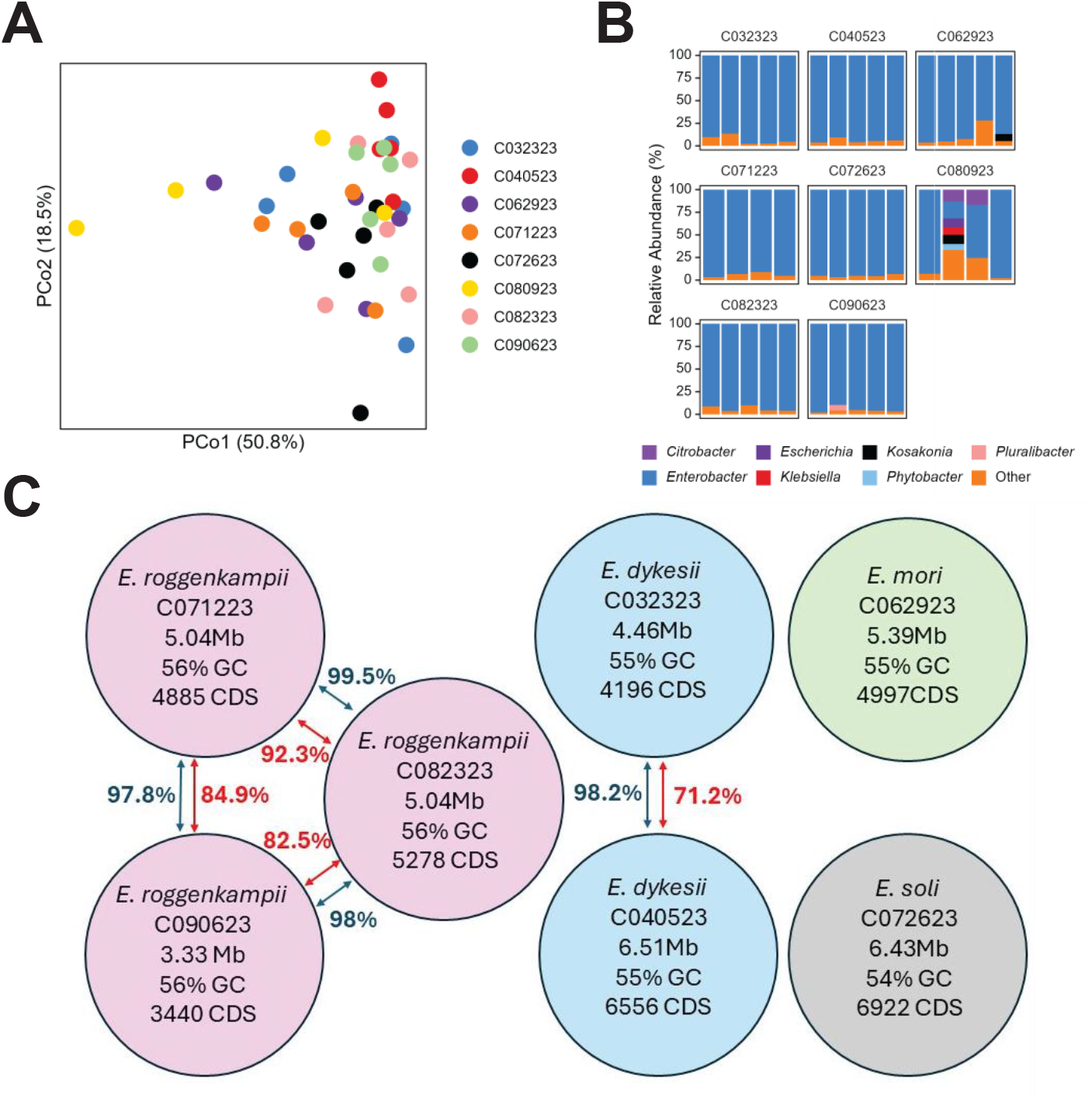
Metagenomic taxonomic composition of mass-reared medfly males and genomes. (A) Principal coordinates analysis of Bray-Curtis dissimilarities and (B) genus level taxonomy of metagenomic reads classified with Kraken2. (C) Co-assembled metagenome assembled genomes indicated no fewer than four *Enterobacter* species and seven *Enterobacter* strains. ANI > 95% reported with blue arrows and synteny (APSS) between species reported in red. Full MAG diagnostics, ANI, and APSS scores and statistics are provided in Supplemental Tables 8-10.

## Discussion

Using near full-length 16S sequencing and co-assembled MAGs through shotgun metagenomics sequencing, we demonstrate that mass-reared medfly males possess compositionally variable microbiomes across a nine-month timeframe. These approaches illuminate differences not previously observed with these same specimens using the V4 region [20]. These results were anticipated, as we previously observed that *in silico* comparisons between full-length 16S and V4 (and V3V4) fragments failed to resolve *Enterobacter* isolates experimentally introduced into medfly [21]. Our data support the notion that long-read 16S sequencing improves resolution and insight into taxon-specific population and community dynamics. This study also exemplifies that a deeper interrogation of insect gut microbiomes using long-read amplicon or whole-genome shotgun metagenomics can reveal greater microbial biodiversity even in ‘simple’ microbial communities.

Whether sequencing longer fragments dramatically improves insight into bacterial community ecology compared to shorter amplicons varies between systems, environmental conditions, and research objectives. A separate study of insects (bees) using full-length 16S sequences also noted improved taxonomic resolution compared to V4 [64], but did not indicate marked differences in dissimilarity metrics. In contrast, in a study of Queensland fruit fly larvae which were dominated by *Asaia* (Acetobacteraceae), there was little discordance between full-length 16S and V4 subregion on statistical signal in the dataset [65]. In other cases, full-length 16S may not provide the resolution needed to parse species or strains, but sequences spanning the length of 16S-ITS-23S can allow for greater strain-level resolution [66]. With improvements to operon database and taxonomic classification [67] along the dramatic increase in amplicon reads obtained with PacBio Kinnex concatenation and newer HiFi sequencing chemistries [22], full operon sequence analysis becomes both economically and technically feasible at a cost similar to, or lower than traditional short read amplicon sequencing depending on planforms used. Some ASVs are certainly arising from one organism being split into multiple sequence variants due to intragenomic copy variation within different isolates or strains, which can inflate biodiversity metrics [68-70], so this may be compounded with even longer operons. At the very least, the full-length 16S sequencing on a PacBio platform has the potential to be accessible and improve resolution. In our case, initially unremarkable results obtained with short-read sequencing [20] were revealed to have a hidden biodiversity through both full-length 16S and shotgun metagenomics, raising new questions related to bacterial establishment, populationdynamics, microbial competition, ecological distributions, and function.

Inferred MAGs largely supported the full-length 16S amplicon results, with different species and strains occurring between the different cohorts. Taxonomic inferences with shotgun metagenomic data were less informative in comparison to the genome-scaled assemblies. This may be due to limitations with species-level classification through tradeoffs between classification confidence and sensitivity (i.e., high confidence thresholds improve precision but reduce the classification rate [71, 72], the composition and quality of reference databases [71], sequence similarity among taxa leading to false positives and negatives [73], or misclassification rates varying substantially by species [74, 75]. It is also important to acknowledge that the sample size is much lower than in the amplicon set of samples, which limits statistical power. A caveat with these data is while they support the long-read data they do not necessarily validate our results.

Bacteria have been shown to mediate tephritid fruit fly development and sexual performance [9, 76], where some bacterial species and strains can impart marked changes to fly physiology and mate selection [77-79]. Tephritid gut systems are frequently populated by Gammaproteobacteria and more specifically, bacteria in the Enterobacteriaceae [9]. Fruit fly gut microbiomes can vary between individual species feeding on distinct fruit [80], which suggests the potential for some environmental acquisition. For medfly, *Klebsiella* and *Enterobacter* are often detected in both culture- and sequencing-based studies in both colony and wild populations [81-85], but how many different species or strains are within a population is still undetermined. Using both full-length sequencing and shotgun metagenomics indicates that there are multiple species and strains associated with medfly, even from the same rearing environment. Currently, we cannot pinpoint the drivers of this variation. One possibility is that there is a carry-over from the larval rearing. Alternatively, there is a rapid depletion and re-population of bacterial titers (from ∼10^2^ to ∼10^7^ CFUs mL^-1^) in the medfly gut following adult emergence from pupae [21, 86], so this may also provide avenues for introduction through food, water, or cage.

Irradiation-induced sterilization was a component of our initial study, which can damage the digestive system and have adverse fitness consequences [87]. Other studies had demonstrated impacts of irradiation on the gut microbiome [79, 88, 89], but outside of one cohort we did not see an overarching effect present across all samples. The differences between our observations and these other studies may be due to depauperate biodiversity above the genus level.

Alternatively, there may have been a lack of opportunistic microorganisms, so there were no large-scale changes in community composition. The level of irradiation used for sterilizing fruit flies (250 gray) are several orders of magnitude below those that would directly affect the bacteria, so we expect potential changes via irradiation would most likely be host-mediated.

Community modules highlighted co-occurrence of ASVs in the medfly microbiome. The most abundant module was present across all cohorts which suggests that while there is variability between cohorts, the most prevalent taxa are linked with each other. Modules were shared across different cohorts, which suggests that the most dominant ASVs may comprise core components of the medfly microbiome but may fluctuate depending upon individual conditions. At least two of these modules correlated with fly mortality, but we caution extrapolation and/or generating causal inferences of these relationships. The number of observations for this comparison were low (n=16) and these trends need support with manipulative experiments.

Our study provides novel insights into the medfly gut microbiome and demonstrates greater bacterial taxonomic complexity in what initially appeared to be unremarkable samples. Full-length 16S sequencing can provide a bridge between short-read and shotgun metagenomics to identify variation to be further interrogated through shotgun metagenomics and MAG assemblies. We anticipate full-length sequences would improve microbiome resolution of other herbivorous insects and systems that have frequent bacterial association within the Enterobacteriaceae.

## Supporting information

Combined Supplements

## Acknowledgements

We thank Renee Corpuz, Angela Kauwe, Tyler Simmonds, and Mikinley Weaver for technical assistance. The findings and conclusions in this publication are those of the authors and should not be construed to represent any official USDA or U.S. Government determination or policy. Mention of trade names or commercial products in this publication is solely for the purpose of providing specific information and does not imply recommendation or endorsement by the U.S. Department of Agriculture. USDA is an equal opportunity provider and employer. This research used resources provided by the SCINet project and/or the AI Center of Excellence of the USDA

Agricultural Research Service, ARS project numbers 0201-88888-003-000D and 0201-88888-002-000D. This research was supported by the U.S. Department of Agriculture, Agricultural Research Service appropriated project “Advancing Molecular Pest Management, Diagnostics, and Eradication of Fruit Flies and Invasive Species” (2040-22430-028-000-D).

## Data Availability

Raw sequence data has been submitted to NCBI SRA under the accession numbers PRJNA1246597 and PRJNA1398906. R scripts and data are available for review at: https://figshare.com/s/b61438ce1df284fb57ad.

---

**Supplemental Figure 1:** Sequence distribution and rarefaction curves of amplicon dataset.

**Supplemental Figure 2:** ASV Richness and Shannon metrics between different cohorts. Different letters represent statistically significant differences (p < 0.05).

**Supplemental Figure 3**: Heatmap of ASVs clustered by sample Bray-Curtis dissimilarities.

## Notes

### Competing Interest Statement

The authors have declared no competing interest.

## References

1. Moran NA, Ochman H, Hammer TJ. Evolutionary and ecological consequences of gut microbial communities. Annual review of Ecology and Systematics. 2019

2. Engel P, Moran NA. The gut microbiota of insects - diversity in structure and function. FEMS microbiology reviews. 2013;37:699–735 10.1111/1574-6976.12025

3. Lange C, Boyer S, Bezemer TM et al. Impact of intraspecific variation in insect microbiomes on host phenotype and evolution. The ISME Journal. 2023;17:1798–807 10.1038/s41396-023-01500-2

4. Shao Y, Mason CJ, Felton GW. Toward an integrated understanding of the Lepidoptera microbiome. Annual Review of Entomology. 2024;69:117–37

5. Qadri M, Short S, Gast K et al. Microbiome innovation in agriculture: Development of microbial based tools for insect pest management. Frontiers in Sustainable Food Systems. 2020;4:547751

6. Abellan-Schneyder I, Matchado MS, Reitmeier S et al. Primer, pipelines, parameters: Issues in 16s rRNA gene sequencing. Msphere. 2021;6:10.1128/msphere.01202-20

7. Ghyselinck J, Pfeiffer S, Heylen K et al. The effect of primer choice and short read sequences on the outcome of 16s rrna gene based diversity studies. PloS One. 2013;8:e71360

8. Bukin YS, Galachyants YP, Morozov I et al. The effect of 16S rRNA region choice on bacterial community metabarcoding results. Scientific Data. 2019;6:1–14

9. Raza MF, Yao Z, Bai S et al. Tephritidae fruit fly gut microbiome diversity, function and potential for applications. Bulletin of Entomological Research. 2020;110:423–37

10. Naum M, Brown EW, Mason-Gamer RJ. Is 16s rDNA a reliable phylogenetic marker to characterize relationships below the family level in the Enterobacteriaceae? Journal of Molecular Evolution. 2008;66:630–42 10.1007/s00239-008-9115-3

11. Abbott SL Klebsiella, Enterobacter, Citrobacter, Serratia, Plesiomonas, and other Enterobacteriaceae. In: Versalovic J, Carroll KC, Funke G et al. (eds.). Manual of Clinical Microbiology, 639–57. Retreived from https://onlinelibrary.wiley.com/doi/abs/10.1128/9781555816728.ch37

12. Drancourt M, Bollet C, Carlioz A et al. 16s ribosomal DNA sequence analysis of a large collection of environmental and clinical unidentifiable bacterial isolates. Journal of Clinical Microbiology. 2000;38:3623–30

13. Dauga C. Evolution of the gyrb gene and the molecular phylogeny of enterobacteriaceae: A model molecule for molecular systematic studies. International Journal of Systematic and Evolutionary Microbiology. 2002;52:531–47 10.1099/00207713-52-2-531

14. Greay TL, Gofton AW, Zahedi A et al. Evaluation of 16S next-generation sequencing of hypervariable region 4 in wastewater samples: An unsuitable approach for bacterial enteric pathogen identification. Science of The Total Environment. 2019;670:1111–24 10.1016/j.scitotenv.2019.03.278

15. Gutiérrez-García K, Whitaker MRL, Bustos-Díaz ED et al. Gut microbiomes of cycad-feeding insects tolerant to β-methylamino-l-alanine (bmaa) are rich in siderophore biosynthesis. ISME Communications. 2023;3:122 10.1038/s43705-023-00323-8

16. Meng Y, Li S, Zhang C et al. Strain-level profiling with picodroplet microfluidic cultivation reveals host-specific adaption of honeybee gut symbionts. Microbiome. 2022;10:140

17. Tedersoo L, Albertsen M, Anslan S et al. Perspectives and benefits of high-throughput long-read sequencing in microbial ecology. Applied and Environmental Microbiology. 2021;87:e00626–21

18. Johnson JS, Spakowicz DJ, Hong B-Y et al. Evaluation of 16s rrna gene sequencing for species and strain-level microbiome analysis. Nature Communications. 2019;10:5029 10.1038/s41467-019-13036-1

19. Gehrig JL, Portik DM, Driscoll MD et al. Finding the right fit: Evaluation of short-read and long-read sequencing approaches to maximize the utility of clinical microbiome data. Microbial Genomics. 2022;8:000794

20. Mason CJ. Evaluating impacts of radiation-induced sterilization on the performance and gut microbiome of mass-reared mediterranean fruit fly (ceratitis capitata) in hawai’i. Journal of Economic Entomology. 2024 10.1093/jee/toae173

21. Mason CJ, Nelson RC, Weaver M et al. Assessing the impact of diet formulation and age on targeted bacterial establishment in laboratory and mass-reared Mediterranean fruit fly using full-length 16S rRNA sequencing. Microbiology Spectrum e02881–24

22. Al’Khafaji AM, Smith JT, Garimella KV et al. High-throughput rna isoform sequencing using programmed cdna concatenation. Nature Biotechnology. 2024;42:582–86

23. Jouvenot Y, Obert C, Hale B et al. The use of iconpcr for 16S library preparation improves data quality and workflow. bioRxiv. 2024:2024.12. 18.629279

24. Mason C, Weaver M, Kissinger K et al. Applying pcr cycle autonormalization to improve pacbio full-length 16S rRNA sequencing. bioRxiv. 2026:2026.01. 13.699363

25. Shen W, Le S, Li Y et al. Seqkit: A cross-platform and ultrafast toolkit for fasta/q file manipulation. PloS one. 2016;11:e0163962

26. Martin M. Cutadapt removes adapter sequences from high-throughput sequencing reads. EMBnet journal. 2011;17:10–12

27. Callahan BJ, McMurdie PJ, Rosen MJ et al. Dada2: High-resolution sample inference from illumina amplicon data. Nature Methods. 2016;13:581–83

28. Callahan BJ, Wong J, Heiner C et al. High-throughput amplicon sequencing of the full-length 16S rRNA gene with single-nucleotide resolution. Nucleic Acids Research. 2019;47:e103–e03

29. R: A language and environment for statistical computing [Computer software]. R Foundation for Statistical Computing. http://www.r-project.org/. 2024.

30. Wang Q, Garrity GM, Tiedje JM et al. Naive bayesian classifier for rapid assignment of rRNA sequences into the new bacterial taxonomy. Applied and Environmental Microbiology. 2007;73:5261–67 10.1128/AEM.00062-07

31. Oksanen, J, Simpson, GL, Blanchet, FG et al. Vegan: Community ecology package [Computer software]. Version R package version 2.6-6.1 <https://CRAN.R-project.org/package=vegan>. 2024.

32. Martinez Arbizu, P. Pairwiseadonis: Pairwise multilevel comparison using adonis [Computer software]. Version R package version 0.4.1 <https://github.com/pmartinezarbizu/pairwiseAdonis>. 2017.

33. Kolde, R. Pheatmap: Pretty heatmaps [Computer software]. Version R package version 1.0.12 https://CRAN.R-project.org/package=pheatmap. 2019.

34. Kurtz ZD, Müller CL, Miraldi ER et al. Sparse and compositionally robust inference of microbial ecological networks. PLoS Computational Biology. 2015;11:e1004226

35. Meinshausen N, Bühlmann P. High-dimensional graphs and variable selection with the lasso. 2006

36. Csardi, G, Nepusz, T, Tragg, V et al. Igraph: Network analysis and visualization in r [Computer software]. Version R package version 2.1.4 <https://CRAN.R-project.org/package=igraph>. 2025.

37. Traag VA, Waltman L, Van Eck NJ. From louvain to leiden: Guaranteeing well-connected communities. Scientific Reports. 2019;9:1–12

38. Wickham, H. Functions to plot networks with ggplot2 [Computer software]. Version R package version 0.1.0, commit da9a7cf2fac24a37ef626e6143ee376eebbdf2d8 <https://github.com/briatte/ggnet>. 2025.

39. Kassambara, A. Rstatix: Pipe-friendly framework for basic statistical tests [Computer software]. Version R package version 0.7.2 <https://CRAN.R-project.org/package=rstatix>. 2023.

40. Andrews S Fastqc: A quality control tool for high throughput sequence data [online].

41. Bolger AM, Lohse M, Usadel B. Trimmomatic: A flexible trimmer for illumina sequence data. Bioinformatics. 2014;30:2114–20

42. Krakau S, Straub D, Gourlé H et al. Nf-core/mag: A best-practice pipeline for metagenome hybrid assembly and binning. NAR genomics and bioinformatics. 2022;4:qac007

43. Di Tommaso P, Chatzou M, Floden EW et al. Nextflow enables reproducible computational workflows. Nature Biotechnology. 2017;35:316–19

44. Kurtzer GM, Sochat V, Bauer MW. Singularity: Scientific containers for mobility of compute. PloS one. 2017;12:e0177459

45. Chen S, Zhou Y, Chen Y et al. Fastp: An ultra-fast all-in-one fastq preprocessor. Bioinformatics. 2018;34:i884–i90

46. Langmead B, Salzberg SL. Fast gapped-read alignment with bowtie 2. Nature methods. 2012;9:357–57

47. Li D, Liu C-M, Luo R et al. Megahit: An ultra-fast single-node solution for large and complex metagenomics assembly via succinct de bruijn graph. Bioinformatics. 2015;31:1674–76

48. Gurevich A, Saveliev V, Vyahhi N et al. Quast: Quality assessment tool for genome assemblies. Bioinformatics. 2013;29:1072–75

49. Wu Y-W, Simmons BA, Singer SW. Maxbin 2.0: An automated binning algorithm to recover genomes from multiple metagenomic datasets. Bioinformatics. 2016;32:605–07

50. Kang DD, Li F, Kirton E et al. Metabat 2: An adaptive binning algorithm for robust and efficient genome reconstruction from metagenome assemblies. PeerJ. 2019;7:e7359

51. Alneberg J, Bjarnason BS, de Bruijn I et al. Concoct: Clustering contigs on coverage and composition. arXiv preprint arXiv:13124038. 2013

52. Sieber CM, Probst AJ, Sharrar A et al. Recovery of genomes from metagenomes via a dereplication, aggregation and scoring strategy. Nature microbiology. 2018;3:836–43

53. Simão FA, Waterhouse RM, Ioannidis P et al. Busco: Assessing genome assembly and annotation completeness with single-copy orthologs. Bioinformatics. 2015;31:3210–12

54. Seppey M, Manni M, Zdobnov EM Busco: Assessing genome assembly and annotation completeness. Gene prediction: Methods and protocols, Springer. 227–45

55. Kriventseva EV, Kuznetsov D, Tegenfeldt F et al. Orthodb v10: Sampling the diversity of animal, plant, fungal, protist, bacterial and viral genomes for evolutionary and functional annotations of orthologs. Nucleic acids research. 2019;47:D807–D11

56. Chklovski A, Parks DH, Woodcroft BJ et al. Checkm2: A rapid, scalable and accurate tool for assessing microbial genome quality using machine learning. Nature methods. 2023;20:1203–12

57. Chaumeil P-A, Mussig AJ, Hugenholtz P et al. Gtdb-tk v2: Memory friendly classification with the genome taxonomy database. Bioinformatics. 2022;38:5315–16

58. Jain C, Rodriguez-R LM, Phillippy AM et al. High throughput ani analysis of 90k prokaryotic genomes reveals clear species boundaries. Nature communications. 2018;9:5114

59. Enav H, Paz I, Ley RE. Strain tracking in complex microbiomes using synteny analysis reveals per-species modes of evolution. Nature Biotechnology. 2025;43:773–83

60. Wood DE, Lu J, Langmead B. Improved metagenomic analysis with Kraken 2. Genome biology. 2019;20:257

61. Lu J, Breitwieser FP, Thielen P et al. Bracken: Estimating species abundance in metagenomics data. PeerJ Computer Science. 2017;3:e104

62. Paulson JN, Pop M, Bravo HC. Metagenomeseq: Statistical analysis for sparse high-throughput sequencing. Bioconductor package. 2013;1:191

63. McMurdie PJ, Holmes S. Phyloseq: An r package for reproducible interactive analysis and graphics of microbiome census data. PloS one. 2013;8:e61217

64. Handy MY, Sbardellati DL, Yu M et al. Incipiently social carpenter bees (xylocopa) host distinctive gut bacterial communities and display geographical structure as revealed by full-length pacbio 16s rRNA sequencing. Molecular Ecology. 2023;32:1530–43 10.1111/mec.16736

65. Deutscher AT, Burke CM, Darling AE et al. Near full-length 16s rRNA gene next-generation sequencing revealed asaia as a common midgut bacterium of wild and domesticated queensland fruit fly larvae. Microbiome. 2018;6:1–22

66. Srinivas M, Walsh CJ, Crispie F et al. Evaluating the efficiency of 16S-ITS-23S operon sequencing for species level resolution in microbial communities. Scientific Reports. 2025;15:2822 10.1038/s41598-024-83410-7

67. Walsh CJ, Srinivas M, Stinear TP et al. Grond: A quality-checked and publicly available database of full-length 16S-ITS-23S rRNA operon sequences. Microbial Genomics. 2024;10:001255

68. Schloss PD. Amplicon sequence variants artificially split bacterial genomes into separate clusters. Msphere. 2021;6:10.1128/msphere.00191-21

69. Earl JP, Adappa ND, Krol J et al. Species-level bacterial community profiling of the healthy sinonasal microbiome using pacific biosciences sequencing of full-length 16S rRNA genes. Microbiome. 2018;6:1–26

70. Sun D-L, Jiang X, Wu QL et al. Intragenomic heterogeneity of 6S rRNA genes causes overestimation of prokaryotic diversity. Applied and environmental microbiology. 2013;79:5962–69

71. Liu Y, Ghaffari MH, Ma T et al. Impact of database choice and confidence score on the performance of taxonomic classification using kraken2. aBIOTECH. 2024;5:465–75

72. Burks DJ, Pusadkar V, Azad RK. Posmm: An efficient alignment-free metagenomic profiler that complements alignment-based profiling. Environmental Microbiome. 2023;18:16

73. Szóstak N, Szymanek A, Havránek J et al. The standardisation of the approach to metagenomic human gut analysis: From sample collection to microbiome profiling. Scientific Reports. 2022;12:8470

74. Chen X, Yin X, Xu X et al. Species-resolved profiling of antibiotic resistance genes in complex metagenomes through long-read overlapping with argo. Nature Communications. 2025;16:1744

75. Govender KN, Eyre DW. Benchmarking taxonomic classifiers with illumina and nanopore sequence data for clinical metagenomic diagnostic applications. Microbial Genomics. 2022;8:000886

76. Deutscher AT, Chapman TA, Shuttleworth LA et al. Tephritid-microbial interactions to enhance fruit fly performance in sterile insect technique programs. BMC Microbiol. 2019;19:287 10.1186/s12866-019-1650-0

77. Behar A, Yuval B, Jurkevitch E. Gut bacterial communities in the mediterranean fruit fly (Ceratitis capitata) and their impact on host longevity. Journal of Insect Physiology. 2008;54:1377–83 10.1016/j.jinsphys.2008.07.011

78. Ben Ami E, Yuval B, Jurkevitch E. Manipulation of the microbiota of mass-reared mediterranean fruit flies Ceratitis capitata (Diptera: Tephritidae) improves sterile male sexual performance. The ISME journal. 2010;4:28–37

79. Cai Z, Yao Z, Li Y et al. Intestinal probiotics restore the ecological fitness decline of Bactrocera dorsalis by irradiation. Evolutionary applications. 2018;11:1946–63

80. Kempraj V, Auth J, Cha DH et al. Impact of larval food source on the stability of the Bactrocera dorsalis microbiome. Microbial Ecology. 2024;87:46

81. Darrington M, Leftwich PT, Holmes NA et al. Characterisation of the symbionts in the mediterranean fruit fly gut. Microbial Genomics. 2022;8:000801

82. Arias MB, Hartle-Mougiou K, Taboada S et al. Unveiling biogeographical patterns in the worldwide distributed Ceratitis capitata (medfly) using population genomics and microbiome composition. Molecular Ecology. 2022;31:4866–83

83. Bel Mokhtar N, Catala-Oltra M, Stathopoulou P et al. Dynamics of the gut bacteriome during a laboratory adaptation process of the mediterranean fruit fly, ceratitis capitata. Front Microbiol. 2022;13:919760 10.3389/fmicb.2022.919760

84. Nikolouli K, Augustinos AA, Stathopoulou P et al. Genetic structure and symbiotic profile of worldwide natural populations of the mediterranean fruit fly, ceratitis capitata. BMC genetics. 2020;21:1–13

85. Aoki S, Weaver M, Simmonds TJ et al. Effects of species and sex on the gut microbiome of four laboratory-reared fruit fly lines (diptera: Tephritidae) using full-length 16s rrna pacbio kinnex sequencing. BMC Microbiology. 2025;25:455

86. Mason CJ, Auth J, Geib SM. Gut bacterial population and community dynamics following adult emergence in pest tephritid fruit flies. Scientific Reports. 2023;13:13723 10.1038/s41598-023-40562-2

87. Lauzon CR, Potter SE. Description of the irradiated and nonirradiated midgut of ceratitis capitata wiedemann (diptera: Tephritidae) and anastrepha ludens loew (diptera: Tephritidae) used for sterile insect technique. Journal of Pest Science. 2012;85:217–26 10.1007/s10340-011-0410-1

88. Woruba DN, Morrow JL, Reynolds OL et al. Diet and irradiation effects on the bacterial community composition and structure in the gut of domesticated teneral and mature queensland fruit fly, Bactrocera tryoni (Diptera: Tephritidae). BMC Microbiology. 2019;19:1–13 10.1186/s12866-019-1649-6

89. Stathopoulou P, Asimakis ED, Khan M et al. Irradiation effect on the structure of bacterial communities associated with the oriental fruit fly, Bactrocera dorsalis. Entomologia Experimentalis et Applicata. 2019;167:209–19

